# *Plasmodium* GPI-Anchored Micronemal Antigen is essential for parasite transmission through the mosquito host

**DOI:** 10.1101/2022.02.24.481744

**Authors:** Charlie Jennison, Janna M. Gibson, Nina Hertoghs, Dorender A. Dankwa, Sudhir Kumar, Biley A. Abatiyow, Myo Naung, Nana K. Minkah, Kristian E. Swearingen, Robert L. Moritz, Alyssa. E. Barry, Stefan H. I. Kappe, Ashley M. Vaughan

## Abstract

The complex life cycle of *Plasmodium* parasites, the eukaryotic pathogens that cause malaria, features three distinct invasive forms tailored specifically to the equally distinct host environment they must navigate and invade for progression of the life cycle. One conserved feature of all these invasive forms is the presence of micronemes, apically oriented secretory organelles involved in egress, motility, adhesion and invasion. Micronemes are tailored to their specific host environment and feature stage specific contents. Here we investigate the role of GPI-anchored micronemal antigen (GAMA), which shows a micronemal localization in all zoite forms of the rodent infecting species *Plasmodium berghei*. While GAMA is dispensable during asexual blood stages, GAMA knock out parasites are severely defective for invasion of the mosquito midgut, resulting in reduced numbers of oocysts. Once formed, oocysts develop normally, however sporozoites are unable to egress and these sporozoites exhibit defective motility. Epitope-tagging of GAMA revealed tight temporal expression late during sporogony and showed that GAMA is shed during sporozoite gliding motility in a similar manner to circumsporozoite protein. Complementation of *P. berghei* knock out parasites with full length *P. falciparum* GAMA partially restored infectivity to mosquitoes, indicating a conservation of function across *Plasmodium* species. A suite of parasites with GAMA expressed under the promoters of the known ookinete-to-sporozoite stage-specific genes: CTRP, CAP380 and TRAP, further confirmed the involvement of GAMA in midgut infection, motility and infection of the mammalian host and revealed a lethal consequence to overexpression of GAMA during oocyst development. Combined, the research suggest that GAMA plays independent roles in sporozoite motility, egress and invasion, possibly implicating GAMA as a regulator of microneme function.

**AUTHOR SUMMARY:** Malaria remains a major source of morbidity and mortality across the globe. Completion of a complex life cycle between vertebrates and mosquitoes is required for the maintenance of parasite populations and the persistence of malaria disease and death. Three invasive forms across the complex lifecycle of the parasite must successfully egress and invade specific cell types within the vertebrate and mosquito hosts to maintain parasite populations and consequently disease and suffering. A conserved feature of all invasive forms are the micronemes, apically oriented secretory organelles which contain proteins required for motility, egress and invasion. Few proteins are expressed in the micronemes of all three invasive forms. One such protein is GPI-anchored micronemal antigen (GAMA). Here we reveal that GAMA is required for the invasion of the mosquito midgut, egress of sporozoites from oocysts and invasion of the vertebrate host. Our finding indicate that while GAMA is essential for sporozoite motility, the defects in oocyst egress and hepatocyte invasion occur independently of the motility defect, implicating the requirement of GAMA in all three processes.

## INTRODUCTION

*Plasmodium* parasites exert a huge morbidity, mortality and economic burden on primarily disadvantaged populations across the globe (1). The complex life cycle of the parasite encompasses three invasive forms, the ookinete, the sporozoite and the merozoite. Within their human or vertebrate secondary host, invasion of the hepatocyte by the sporozoite and the red blood cell by the merozoite is essential for these obligate intracellular parasite life stages. Invasion relies on an array of proteins tailored for invasion of the relevant host cell type, proteins housed in secretory organelles, primarily the micronemes and rhoptries, which are located toward the apical tip of the invasive zoites (2–4). Rhoptries are absent from the mosquito-invasive form, the ookinete, as they house proteins required specifically for host cell invasion; while the ookinete must “invade” the mosquito midgut, this process could also be viewed as a traversal event through the midgut epithelium, as differentiation and the growth of oocysts occurs extracellularly between the basal lamina (an extracellular matrix supporting and surrounding the midgut) and the epithelium cells of the midgut (5–7). Micronemes are therefore the most important conserved feature across these motile forms, yet relatively few *Plasmodium* micronemal proteins present in all three invasion stages are characterized and little is known about microneme formation and secretion (8–10).

One understudied micronemal protein, implicated in the progression of all invasive life cycle forms is GPI anchored micronemal antigen (GAMA). *GAMA* was deleted in a reverse genetic screen of proteins selected for their ookinete expression and suspected secretion, first described as **P**utative **S**ecreted **O**okinete **P**rotein ***9*** (PSOP9), thereby revealing non-essentiality in the asexual blood stages in the rodent malaria parasite *P. berghei* (11). Δ*psop9/GAMA* parasites exhibited no phenotypic difference to wild type parasites in the blood stage, but at the ookinete stage they were ∼80% defective for invasion of the mosquito midgut. Δ*psop9/GAMA* sporozoites were also unable to infect mosquito salivary glands or establish blood stage infection in mice when sporozoites derived from the oocyst were injected intravenously. The nature of these ookinete and sporozoite defects however remains undetermined.

In the blood stages of the human malaria-causing parasite *P. falciparum*, GAMA (then PF08_0008, reannotated as PF3D7_0828800) was identified in a screen for GPI anchored proteins (12). Immunoprecipitation of GAMA from parasites metabolically labelled with the radioactive GPI precursors [^3^H]mannose and [^3^H]glucosamine further confirmed the *in silico* prediction of a GPI anchor at the C-terminus (13). GAMA was localized to merozoites within mature schizonts, colocalizing with AMA1 thereby indicating an apical and likely micronemal localization, which relocates to either a cap or peripheral localization upon merozoite egress from schizonts (13). Immunoelectron microscopy further confirmed the presence of GAMA in some micronemes (14). In both *P. falciparum* studies of this protein, GAMA was shown to bind red blood cells and Arumugam *et al*. found that antibodies raised against GAMA inhibited blood stage parasite growth in a sialic acid independent manner (13,14). Failed knock out attempts and an absence of mutations in a genome-wide *PiggyBac* transposon mutagenesis screen indicate GAMA may be essential for *P. falciparum* asexual blood stage replication (14,15). The aforementioned findings all point to a role of GAMA in *P. falciparum* during erythrocyte invasion.

Our data support additional roles for GAMA during mosquito stage development where we find it is required not only for ookinete midgut invasion but also for sporozoite egress from the oocyst, sporozoite motility and sporozoite invasion of the mammalian host. Data from our mutant parasites reveal that GAMA plays independent roles in these processes and we propose a model in which GAMA plays an important role in microneme function, with its absence resulting in a cascade of microneme related defects.

## RESULTS

### Sequence analysis of GAMA reveals high conservation across *Plasmodium* species

*P. berghei* GAMA consists of 625 aa with a predicted molecular mass of ∼71 kDa, (signal peptide cleavage results in a 69 kDa mass). In *P. falciparum*, two additional asparagine rich repeat regions result in a larger 738 aa protein (predicted molecular weight ∼85 kDa, 83 kDa after cleavage of the signal peptide). For *P. berghei*, SignalP-5.0 predicts a signal peptide with a cleavage site at S21/L22 (probability of 0.67, likelihood 98% [indicated in Figure 1A]). For *P. falciparum*, cleavage is predicted at the same locus, A21/L22 (probability 0.64, likelihood 99%).

**Figure 1:**
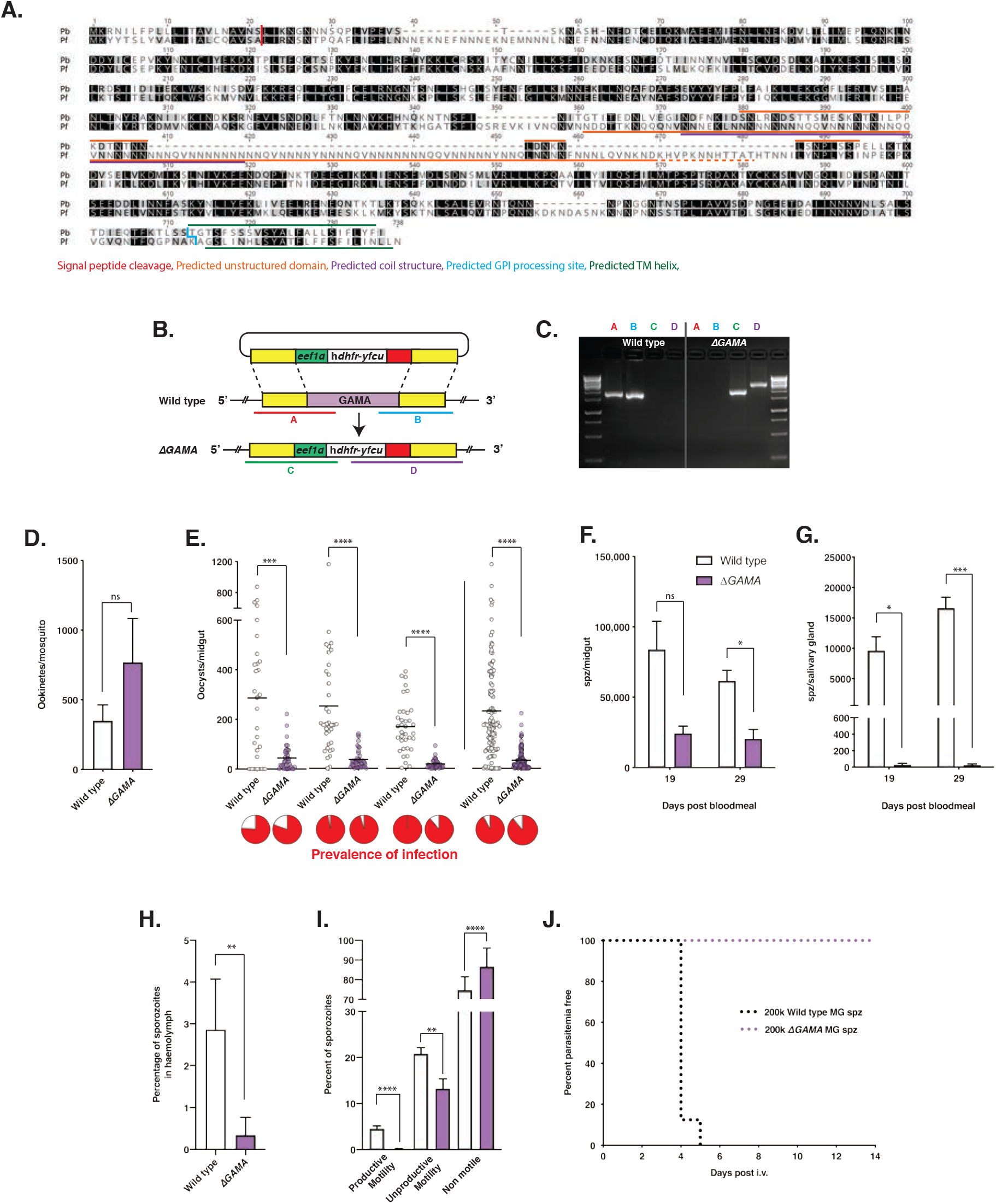
GAMA amino acid alignment and the phenotype of Δ*GAMA* parasites. A. Protein alignment of *P. berghei* and *P. falciparum* GAMA amino acid sequences, showing: predicted signal peptide cleavage site at position 21-22 (red line), predicted unstructured domains (orange line), predicted coil structure in *P. falciparum* (purple line), location of predicted GPI cleavage and attachment (blue line), location of predicted TM helix for anchoring in the endoplasmic reticulum (dark green line). B. Schematic of strategy used to obtain Δ*GAMA* parasites with PCR amplicons used for genotyping annotated A-D. C. Electrophoresis gel confirming disruption of *GAMA*, PCR amplicons from (B.) show above the gel. DNA from wildtype parasites on the left and a clone of Δ*GAMA* on the right. D. Number of ookinetes per mosquito at 22 hours post infectious blood meal, mean and SEM of three biological replicates (paired t-test). Wild type in white, Δ*GAMA* in purple. E. Number of oocysts per midgut, three biological replicates to the left, with all data combined on the right. Horizontal bars indicate mean, prevalence of infection indicated as red portion of pie charts below, (Mann Whitney test performed). F. Number of sporozoites per midgut at two timepoints, mean and SEM of 5 biological replicates (paired t-test). G. Number of sporozoites per mosquito salivary gland at two timepoints, mean and standard deviation of three biological replicates (paired t-test). H. The percentage of haemolymph sporozoites from the total collected from both haemolymph and midguts of mosquitoes infected with wildtype and Δ*GAMA* parasites (t test, p = 0.0077). I. Percentage of wild type and Δ*GAMA* midgut sporozoites that were fully motile, unproductively motile or non-motile, two biological replicates (wild type n = 2687, Δ*GAMA* n = 2046, p < 0.0001, 0.0039 and <0.0001 respectively). J. Time to blood stage patency following intravenous injection with 200,000 oocyst sporozoites, two biological replicates combined, total of eight mice per group.

A sequence alignment of *P. berghei* and *P. falciparum* protein sequences reveals the protein to be highly conserved between these distantly related species. *P. berghei* and *P. falciparum* share 44.8% identity across the full-length sequence, with areas of higher conservation found towards the N-terminus (residues 64-347 = 60% identity) and C-terminus (residues 501-700 = 60% identity) split by a central repeat region of low shared identity (residues 348-500, Fig 1A), with predicted disorderly structure. The C-terminal transmembrane domain is predicted to be cleaved and appended with a GPI anchor (PredGPI, 100% and 99.8% specificity for *P. berghei* and *P. falciparum* respectively, cleavage site indicated in blue on Fig 1A). Other than these, the protein has no predicted functional domains. The predicted disorderly domain in *P. berghei* overlaps with the central asparagine rich repeat region in *P. falciparum* which is predicted to be coiled in structure. Downstream of signal peptide cleavage, *P. falciparum* contains seven cysteine residues prior to the central repeat region and one downstream which are conserved across all *Plasmodium* species investigated (Supp Fig. 1). *P. berghei* and other rodent infective species of *Plasmodium* bear an additional cysteine at position 152 in the alignment (Fig 1, position 170 in Supp Fig. 1). GAMA is present in all *Plasmodium* species and indeed all other intracellular apicomplexans (16).

Based on a dataset of 2668 *P. falciparum* genomes from 15 countries across Asia, Africa, and Papua New Guinea, GAMA was found to be a relatively highly conserved protein compared to other surface-exposed micronemal proteins such as apical membrane antigen 1 (AMA1) (17)and thrombospondin related anonymous protein (TRAP) (Supp Fig 2A) (18,19). Sliding window Tajima’s D analysis did not identify any immune selection hotspots within GAMA across isolates from a broad geographical distribution, however, regions with weak immune selection were detected near the N-terminal of GAMA from Papua New Guinea (n=156, Supp. Fig 2B), and Ghanaian parasite isolates (n=245, Supp. Fig. 2C). In addition, the 3D7 allele is totally absent from field isolates. Taken together these results indicate that GAMA is either highly functionally conserved, thereby restricting sequence diversity, or that GAMA is poorly immunogenic and/or minimally exposed to host immune selection pressures.

For *P. berghei*, binding sites for the ookinete specific transcription factor AP2-O: TAGCTA and TGGCTA, occur 753 (2055bp in *P. falciparum*) and 589 bases upstream of the open reading frame (ORF) respectively, in keeping with the observed expression profile and role in ookinete function (20,21). The binding site for the sporozoite-specific transcription factor AP2-Sp, TGCATGCA, is commonly found in the first 1000 bases upstream of sporozoite expressed genes (22)and we further identified this consensus binding site to be located 1426 upstream of the ORF (1787 in *P. falciparum*), significantly further upstream than other previously investigated sporozoite expressed genes (22). Further sequence analysis indicates that the most upstream expressed sequence tags for *P. berghei* GAMA begin correspondingly distal to the ORF (1140bp) and the 5’UTR of *P. falciparum* is 1728bp in length (PlasmoDB). The long promoter region of GAMA may result from the complex multistage expression profile, thereby requiring multiple transcription factor binding sites. Interestingly, another binding motif, GTGCAC, associated with the transcription factor AP2-D3, found upstream of many schizont expressed genes with functions in merozoite invasion was not found (23), nor were other conserved *cis*-regulatory elements associated with microneme genes: PfM20.1 (ACAACCT) and PfM18.1 (NGGTGCA) (24).

### GAMA is non-essential for *P. berghei* asexual or sexual blood stages

In order to investigate the function of GAMA in *P. berghei*, a gene in marker out (GIMO) strategy was employed (25), whereby a fusion of the positive selectable marker human dihydrofolate reductase (hDHFR) and the negative selectable marker yeast fusion cytosine deaminase/uracil phosphoribosyl-transferase (yFCU) is inserted into the genome via double homologous recombination, thereby removing the GAMA gene and permitting further genetic manipulation (Fig 1B). The parasite line Pb676, a marker-free *P. berghei* parasite expressing GFP under the control of the EF1a promoter, was used as the background parasite line for these transfections and is referred to as wild type from here on (26). Δ*GAMA* parasites were readily obtained, and no blood stage phenotype was observed (Fig. 1C). To investigate the role of GAMA in gametocyte formation and fertilization, mosquitoes were fed on mice infected with either wildtype or Δ*GAMA* parasites at comparable parasitaemia. At 22 hours post infectious blood meal, mosquito midguts were dissected to determine ookinete numbers. Ookinete counts revealed no significant difference between Δ*GAMA* and wildtype ookinete numbers (Fig 1D).

### GAMA is required during ookinete establishment of midgut infection

While the number of ookinetes was not significantly different between Δ*GAMA* and wildtype, the intensity of oocyst infection in the midguts of mosquitoes on day 14 post infectious blood meal was significantly lower in Δ*GAMA* infected mosquitoes, across five biological replicates (Fig 1E; p = 0.0004, p < 0.0001, p < 0.0001, combined data p < 0.0001, additional data in Figure 5E; p < 0.0001, p < 0.0001), indicative of a defect in the ability of ookinetes to invade the midgut epithelium and establish infection. Prevalence of infection between wildtype and Δ*GAMA* infected mosquitoes was not significantly different (Fig. 1E, pie-charts).

### GAMA is not required for sporozoite development but is essential for egress from oocysts and subsequent infection of the salivary glands

Due to the reduced oocyst numbers in Δ*GAMA* infected mosquitoes, there were consequently fewer sporozoites in the midguts of Δ*GAMA* infected mosquitoes at both day 19 (p = 0.03)), and day 29 post blood meal (Fig 1F, p = 0.04). No significant decrease in the number of midgut sporozoites for either wildtype or Δ*GAMA* was observed between days 19-29, however a time course running from day 19-40 revealed a trend of decreasing sporozoite numbers at latter timepoints for wildtype, while midgut sporozoite numbers remained constant over time for Δ*GAMA* (Supp. Fig. 3A). A striking and significant difference in salivary gland sporozoite numbers between wildtype and Δ*GAMA* infected mosquitoes was observed at all timepoints across five replicates (Fig 1G, Paired t-test: day 19; p = 0.013, day 29; p = 0.0007); indeed, no sporozoites were found in the majority of salivary gland dissections for Δ*GAMA* infected mosquitoes, as far out as day 40 post blood meal (Supp. Fig. 3B), with oocysts remaining intact in mosquito midguts, in keeping with previous observations (11). To confirm this defect in sporozoite egress from oocysts, sporozoites were collected from the haemolymph and midguts of the same mosquitoes and the proportion of the combined total present in the haemolymph determined (Fig. 1H). A salivary gland invasion defect would result in a build-up of haemolymph sporozoites which was not observed; indeed, very low numbers of sporozoites were detected and the proportion of sporozoites detected in the haemolymph was eight-fold lower in Δ*GAMA* infected mosquitoes (t test p = 0.0077), confirming a defect in sporozoite egress from the oocyst and a role for GAMA in this process. These results indicate that GAMA is not necessary for the formation of sporozoites but is required for their release from the oocyst, either through direct involvement in egress or by facilitating the function of other egress essential proteins such as ECP1 (27).

### GAMA is essential for the productive motility of sporozoites

While a role for GAMA in the motility of sporozoites has previously been hypothesized (11,28), it has never been experimentally investigated. Midgut sporozoites are known to be much less motile than salivary gland sporozoites. However, as salivary gland sporozoites were unobtainable for Δ*GAMA* parasites and haemolymph sporozoites extremely sparse, midgut sporozoites were the only option for investigating sporozoite motility. Motility in sporozoites is commonly divided into two main categories: productive, unidirectional movement; and unproductive motility, which includes waving from one fixed end, flexing or bending, and back-and-forth patch-gliding, whereby no progress is made (29). To investigate both productive and unproductive motility in a more physiologically relevant environment than the 2D assays often used (imaging of motility or trails on flat glass or plastic surfaces), sporozoites were purified from infected midguts and set in a Matrigel matrix in the presence of bovine serum albumin (BSA) for activation (30,31), with time-lapse footage acquired every second over seven minutes. Motility of each sporozoite was classified into three categories; immotile, unproductive patch-gliding motility (back-and-forth movement of less than one body length), or motile (unidirectional movement of over one body length [Supp. Video 1]). Waving motility was never seen, and flexing/bending, while observed, was extremely rare. As expected, productive motility was low even in wild type sporozoites (112 of 2687, 4.5%), however productive motility in the Δ*GAMA* sporozoites was almost completely absent (p < 0.0001, 3 of 2046, 0.15% [the distance travelled for the three sporozoites just exceeded the one body length required to classify as motile]) (Fig. 1I). Patch gliding was more frequently observed than productive motility for both wild type and Δ*GAMA* sporozoites (21% and 13% respectively), and again this was significantly more common in wild type than Δ*GAMA* parasites (p = 0.004). While these data were generated with a different motility assay, the patch-gliding phenotype observed in our Δ*GAMA* parasites mirror two previous observation of Δ*TRAP* parasites and implicate either a role in motility which manifests with this convergent phenotype, or a role for GAMA in the correct release or localization of TRAP (29,32).

### GAMA is essential for infection of the mammalian host

Sporozoites mature as they transition from oocysts to the salivary glands (33), with midgut sporozoites estimated to be 10,000-fold less infectious than salivary gland sporozoites for the mammalian host (34). Given the oocyst egress defect of Δ*GAMA* sporozoites it was not possible to obtain adequate numbers of salivary gland sporozoites to investigate the role of GAMA in mature sporozoite infection of the mammalian host. Midgut sporozoites were however abundant for both wildtype and Δ*GAMA* parasites and high numbers of these sporozoites were used to investigate sporozoite infectivity. C57BL/6 mice injected intravenously with 200,000 midgut wild type sporozoites (day 21-29 post blood meal) all became blood stage patent by day 6 (Fig 1J & expanded in Fig 5I) whereas 0/21 mice injected with Δ*GAMA* sporozoites developed blood stage infection (Fig. 1J & expanded in 5J). While midgut sporozoites are known to be orders of magnitude less infectious than those that have matured in the salivary glands, these data clearly implicate a role for GAMA either in the establishment of the liver stage – either from a defect during hepatocyte invasion, or as a result of immotility and a failure to traverse through the liver sinusoid and parenchymal cells. Given the accumulated defects up to this point, pinpointing the pre-erythrocytic phenotype with Δ*GAMA* midgut sporozoites is currently not possible.

### The phenotypically silent insertion of an mCherry tag in the C-terminus of GAMA reveals the tightly regulated temporal expression and localization of GAMA across the life cycle

A CRISPR/Cas9 methodology was used to insert an mCherry tag sequence toward the C-terminus of the endogenous GAMA gene five amino acids from the predicted GPI anchoring location (Fig. 2A). Tagged parasites (*GAMA-mCherry*) were readily generated (Fig. 2B) and imaging of mature blood stage schizonts revealed mCherry signal predominantly localized in puncta toward the exterior facing end of merozoites (Fig. 2G, Supp Fig. 4), typical of an apical localization and consistent with previous findings in *P. falciparum* (13,14), indicating protein localization was unaffected by epitope tagging and conserved across species.

**Figure 2:**
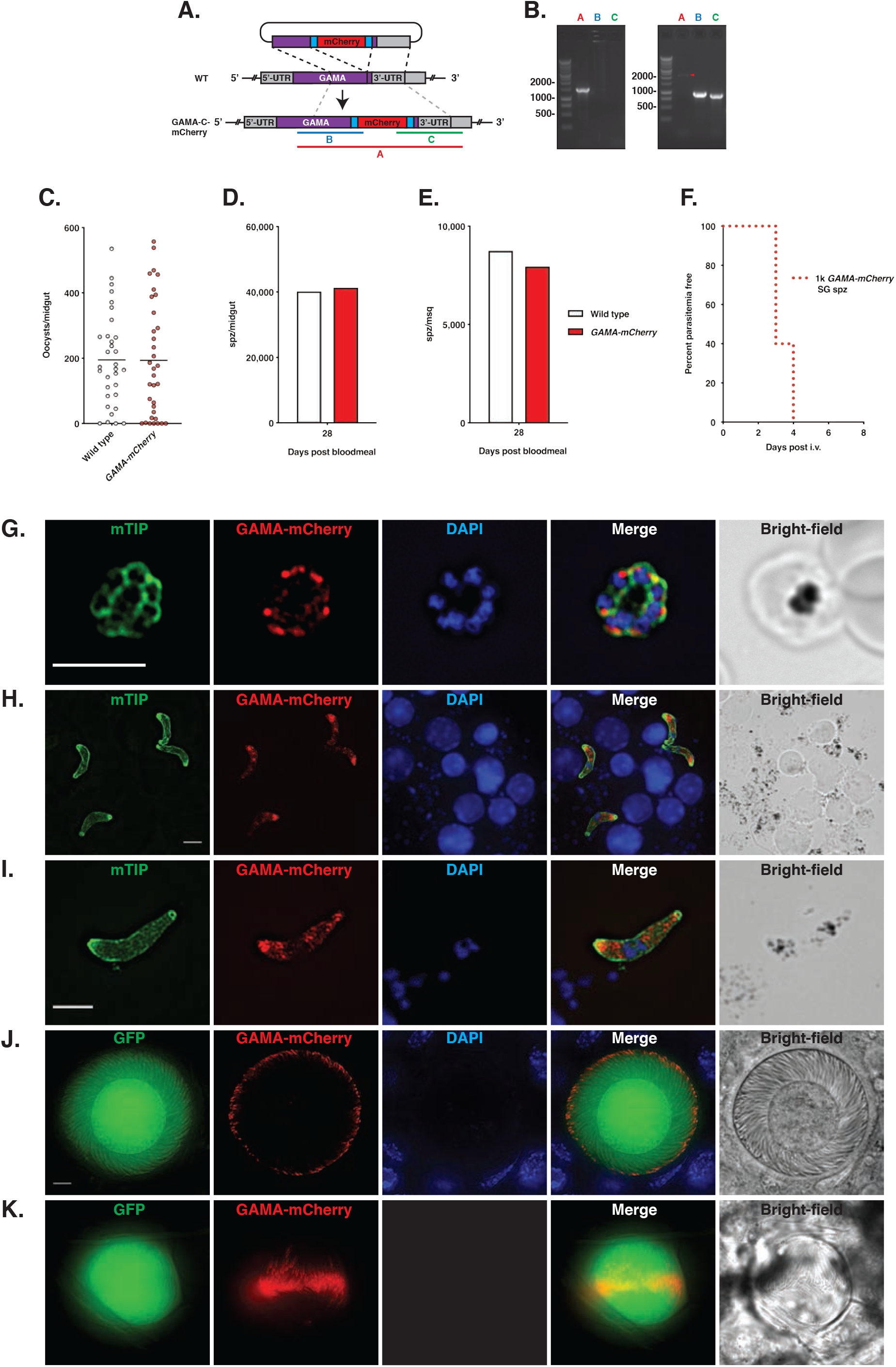
A *P. berghei* parasite with mCherry-tagged GAMA reveals the localization and temporal expression of GAMA across the life cycle. A. Schematic of the tagging strategy for the C terminal tagging of *PbGAMA* with approximate PCR amplicons used for genotyping. B. PCR gel confirming mCherry tagging of GAMA. A (wt = 1501, Tag = 2230), B (wt = N/A, Tag = 947), C (wt = N/A, Tag = 874). Gel on left = wildtype, on right is a representative clone of *Pb-GAMA-C-mCherry*. C. Day 14 oocyst counts for wildtype and *Pb-GAMA-C-mCherry*. D. Midgut sporozoite counts at day 28 post infectious bloodmeal. E. Salivary gland sporozoite numbers at day 28 post infectious bloodmeal. F. Kaplan Meier analysis of five mice injected with 1,000 *Pb-GAMA-C-Cherry* salivary gland sporozoites. G. Immunofluorescence assay of Pb-GAMA-C-mCherry schizont, 1000x magnification, *Pb-GAMA-C-Cherry* schizonts stained with rat α-mCherry (secondary 594, red), rabbit α-*P. yoelii*-MTIP (secondary 647, green), DAPI (blue). Scale bar 5 µm. H. IFA showing large field view, 1000x magnification. *Pb-GAMA-C-mCherry* ookinetes, stained as in (G.). Scale bar 5 µm. I. IFA of *Pb-GAMA-C-mCherry* ookinete, 1000x magnification, staining as in (G.). Scale bar 5 µm. J. Endogenous GFP and mCherry fluorescence in *Pb-GAMA-C-mCherry* oocysts at day 17 post blood meal, showing expression towards the apical tip of budding sporozoites. K. Day 21 oocyst with GAMA-C-mCherry localized in mature sporozoites.

Further phenotypic analysis of *GAMA-mCherry* revealed no detectable changes in the number of oocysts per midgut (Fig. 2C), the number of sporozoites per midgut (Fig. 2D), the number of sporozoites per salivary gland (Fig. 2E), and the infectiousness of these sporozoites to mice (Fig. 2F). In light of the strong phenotypes observed across these life cycle stages in Δ*GAMA* parasites we concluded that our tagging strategy did not interfere with the function of GAMA and therefore used *GAMA-mCherry* to further investigate the localization and temporal expression of GAMA.

*GAMA-mCherry* ookinetes, harvested from mosquitoes 22 hours post infectious blood meal, revealed internal localization of mCherry staining throughout the ookinetes, frequently accumulated towards the wider apical end (as determined by ookinete morphology (Fig. 2H & I)) (35). Oocysts were imaged over the course of their maturation from day seven onwards *ex vivo*. At day seven, using a long exposure time, weak mCherry signal was visible in *GAMA-mCherry* oocysts but not in wild type controls (Supp Fig 4A). No significant increase in mCherry signal was observed until day 11, when a subset of oocysts became brightly mCherry positive (Supp Fig 5B & C). The extremely low-level fluorescence observed in early oocysts is therefore most likely residual mCherry that persists from prior expression during the ookinete stage. A more detailed investigation of *GAMA-mCherry* infected midguts revealed that within the asynchronously matured population of oocysts, it was those that contained developing or mature sporozoites that were mCherry positive (Supp. Fig. 5D) and that mCherry was first localized within the nascent apical tip of developing sporozoites (Fig. 2J, Supp Video 2) (8,36). mCherry localization becomes less restricted to the apical tip as sporozoites matured (Fig 2K & L), replicating the tip-backwards maturation of micronemes within developing sporozoites (8). Within oocysts containing mature sporozoites, mCherry was absent from regions with nuclear staining, corresponding to approximately the midpoint of a sporozoite, indicating the polarized localization of GAMA in oocyst sporozoites (Supp. Fig. 6A). Imaging wet preparations of fresh mosquito midguts 21 days post infectious blood meal revealed GFP negative, mCherry positive regions, corresponding to previously ruptured oocysts (Supp. Fig. 5C) and oocysts were also observed leaking a diffuse cloud of both GFP and mCherry, (Supp. Fig 6B), suggesting that mCherry-GAMA is released during sporozoite egress.

Live-imaging of midgut sporozoites recovered from mechanically ruptured mosquito midguts found *GAMA-mCherry* to be localized within sporozoites, occupying the same space as cytoplasmic GFP, with a somewhat punctate localization (Fig 3A). There was frequently an accumulation of *GAMA-mCherry* towards one end of these sporozoites, however this was inconsistent and may represent the mixed maturity of sporozoites recovered from oocysts (Fig 3A), with recently formed sporozoites showing greater expression toward the apical tip and more mature sporozoites exhibiting a more even distribution throughout. Salivary gland sporozoites, dissected in Schneider’s medium on ice, exhibited a localization of *GAMA-mCherry* similar to that observed in midgut sporozoites, however there was an increase in the abundance of protein (Fig 3B). Using immunofluorescence assay (IFA) detection on mature sporozoites further confirmed the localization seen with endogenous fluorescence (Fig. 3C). Our previous mouse infection experiments with Δ*GAMA* midgut sporozoites indicated an important role of GAMA in the infection of the mammalian host. Thus, to investigate whether GAMA localization was altered upon mosquito injection into the mammalian host, salivary gland sporozoites were incubated for 15 minutes at 37°C in 1% BSA to mimic activation conditions brought on by sporozoite introduction into the dermis during mosquito feeding. Motile sporozoites were imaged gliding on glass coverslips at two second intervals and revealed a similar internal localization to that seen in non-activated salivary gland sporozoites (Fig 3D). The fluorescence intensity of mCherry diminished over time due to photo-bleaching, revealing a greater abundance of protein toward the apical tip. Punctate accumulations of protein could be followed over time in moving sporozoites and did not reveal any clear relocalisation during the process of gliding motility (Fig 3D). The micronemal localization of GAMA (14) and the GPI anchor (12,13) would suggest that GAMA is ultimately incorporated into the parasite membrane. To investigate localization, line-plot pixel intensity analysis across the apical and mid/posterior regions of sporozoites was performed, revealing a concentration of *GAMA-mCherry* inside the apical tip of sporozoites which shifted to a peripheral localisation toward and rear of the sporozoite, suggesting that GAMA is incorporated into the plasma membrane (Fig 3E). Our motility assay had previously revealed an essential role for GAMA in the productive motility of midgut sporozoites. Sporozoite surface proteins can be shed from the sporozoite surface during gliding motility, as is the case with circumsporozoite protein (CSP) and TRAP (30,37), or not, as in the case of sporozoite surface protein 3 (SSP3) (38). To investigate whether GAMA is shed from the sporozoite surface during gliding motility, wild type and *GAMA-mCherry* salivary gland sporozoites were incubated in media containing BSA at 37°C for 45 minutes in order to facilitate motility and the deposition of trails on FBS coated chamber slides. Trails were then stained with antibodies to CSP and mCherry, which clearly revealed the presence of mCherry in the trails of *GAMA-mCherry* parasites (Fig. 3F). In summary, these data show GAMA to be concentrated at the apical tip of sporozoites where it is then translocated to the sporozoite surface and shed during the process of gliding motility. This further suggested that GAMA is involved in sporozoite gliding motility and likely essential for the passage of the sporozoite from the mosquito bite site to its hepatocyte target.

**Figure 3:**
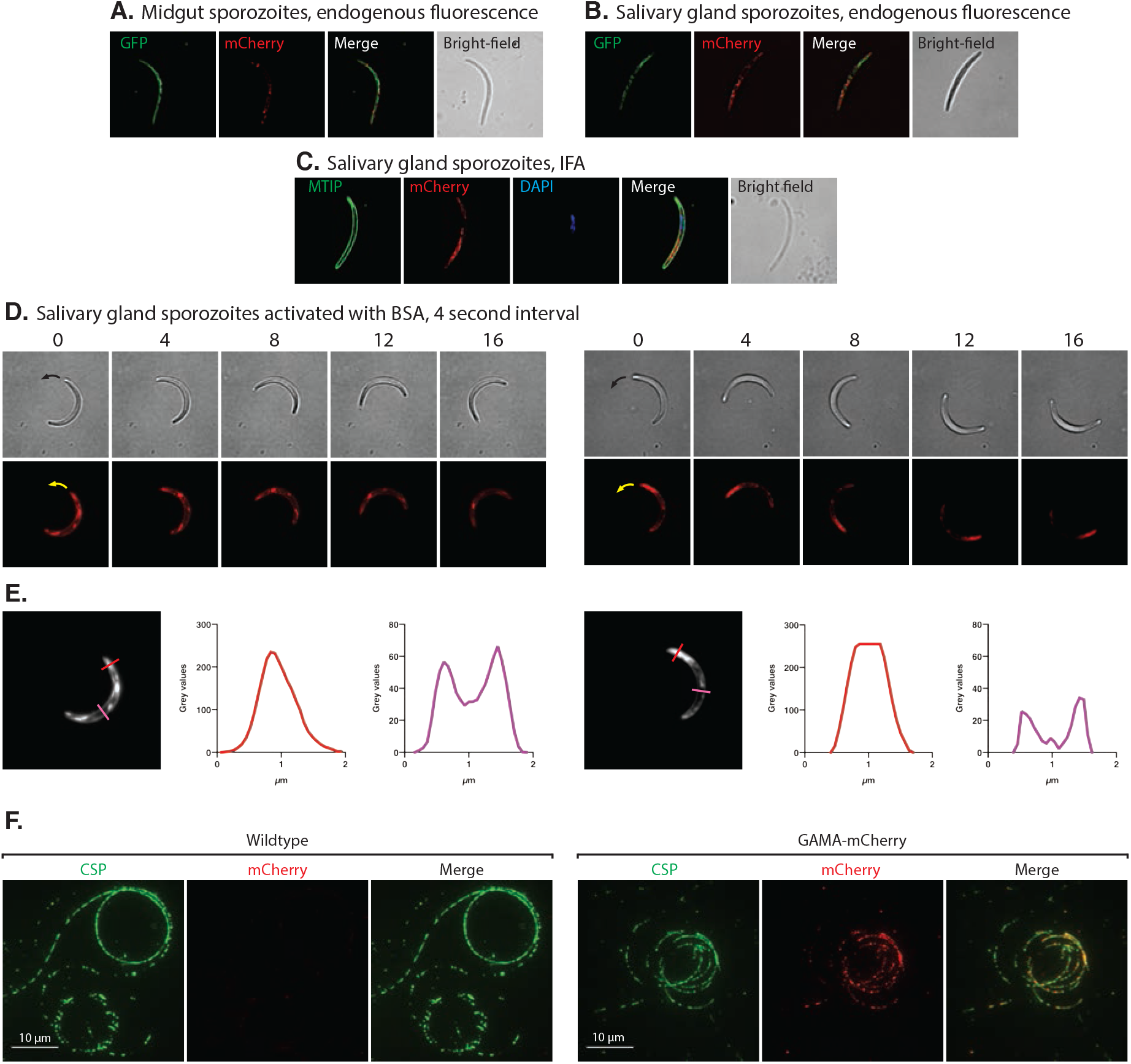
Localisation of GAMA-mCherry in sporozoites. A. Live midgut sporozoites imaged immediately post extraction from midguts, exhibiting low level, punctate mCherry signal within the sporozoite. B. Live salivary gland sporozoites imaged immediately following extraction from salivary glands, showing intense mCherry signal within the sporozoite. C. IFA on paraformaldehyde fixed and permeabilized GAMA-mCherry salivary gland sporozoites stained with rabbit α-MTIP, rat α-mCherry and DAPI. D. Two time-lapse series of salivary gland sporozoites on a glass coverslip, activated with 1% BSA. Frames captured every 4 seconds. mCherry is unevenly distributed within the sporozoite, in some cases with an accumulation toward the apical end of the sporozoite. The requirement for a 0.2 second exposure time results in a slight blurring in the fluorescence images and mCherry signal bleaches over time. E. Intensity/localization of GAMA-mCherry was determined with line-plots of pixel intensity across apical and mid/posterior regions of activated sporozoites. Coloured lines indicate the transect measured, grey value plots correspond to colour matched lines. F. Salivary gland sporozoite trails contain GAMA and CSP. Trails were stained with rat α-mCherry and labelled α-CSP antibody. Microscope settings were consistent between parasite lines.

### Complementation of Δ*GAMA* parasites with *P. falciparum GAMA* results in a stage-specific restoration of function during mosquito infection

Previous work investigating the role of GAMA in *P. falciparum* found the gene refractory to deletion (14)and more recently, *GAMA* was defined as non-mutable in a high throughput *piggyBac* transposon mutagenesis screen, with no insertions observed at a possible 45 sites within *GAMA* (15), further suggesting an essential role in the blood stage of *P. falciparum*. The N- and C-termini of *P. berghei* and *P. falciparum* GAMA are highly conserved (Fig 1A, Supp Fig. 1), and the expression profile is similar (Fig 4I). Therefore, to investigate the role of *P. falciparum* GAMA in non-erythrocytic stages of the lifecycle we complemented our *Pb*Δ*GAMA* parasite with the full-length open reading frame of *P. falciparum GAMA*, generating *Pb*^*PfGAMA*^ parasites to ascertain if there was conservation of function between species adequate to rescue any of the defects observed in Δ*GAMA* parasites (Fig 4A & B).

**Figure 4:**
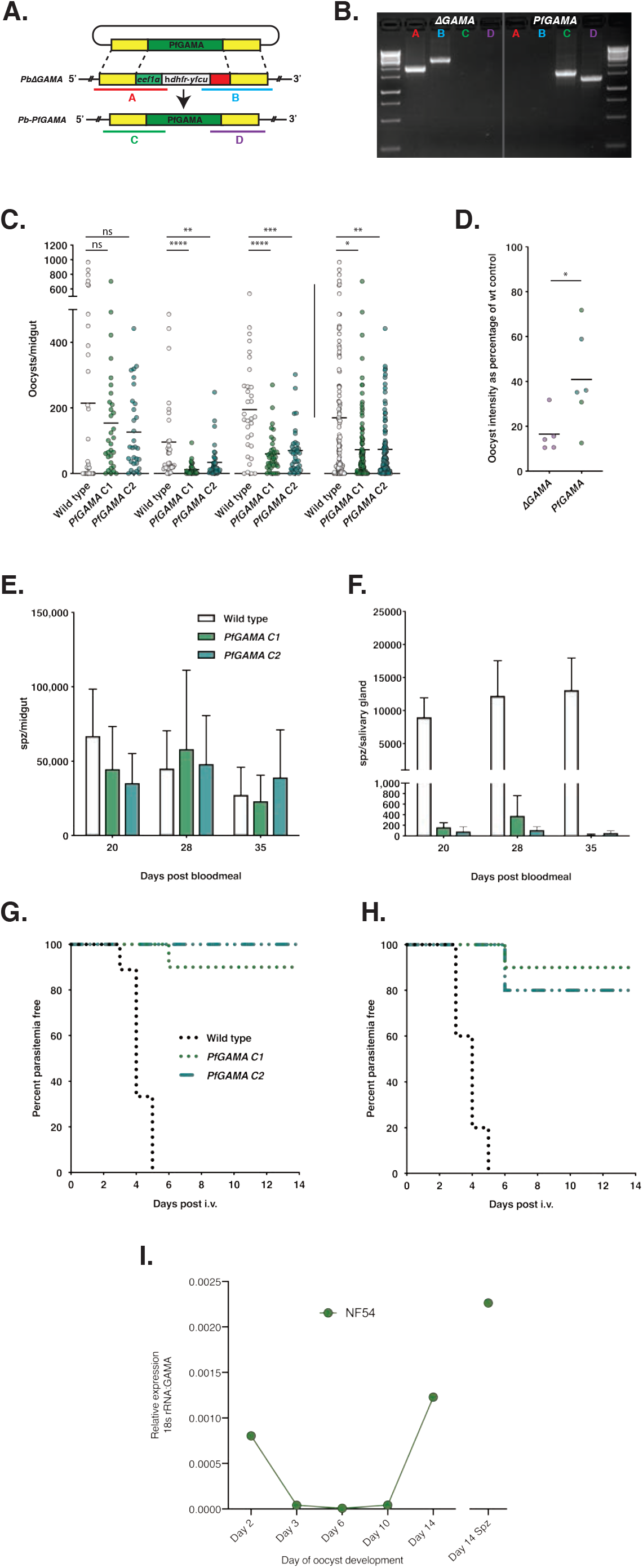
Complementation with *P. falciparum* GAMA partially restores Δ*GAMA* defects across the life cycle. A. Schematic showing the Gene-In-Marker-Out strategy used to complement the Δ*GAMA* parasite with *PfGAMA* with approximate PCR amplicons used for genotyping. B. PCR gel confirming replacement of the hDHFR/yFCU cassette with the GAMA gene from *P. falciparum*. C. Oocyst counts of mosquito midguts 14 days post infectious blood meal, three biological replicates. To the right of the vertical line are all three replicates combined. Mean shown, (Mann-Whitney test, p > 0.05 = ns, p < 0.05 = *, p < 0.01 = **, p < 0.001 = ***, p < 0.0001 = ****). D. Percent reduction in oocyst intensity when compared with respective control wildtype infections for Δ*GAMA* and *PfGAMA* parasites, mean shown, (one tailed Mann-Whitney, *p = 0.026) E. Midgut sporozoites per mosquito at three timepoints, mean and SEM of three biological replicates. F. Salivary gland sporozoites per mosquito at three timepoints, mean and SEM of three biological replicates. G. Kaplan Meier survival curve showing time to blood stage patency following intravenous injection of 200,000 midgut sporozoites per C57BL/6 mouse, two biological replicates combined, n = (wt = 9, clone 1 = 10, clone 2 = 10). Twenty fields of a Giemsa-stained thin smear checked daily. H. Kaplan Meier survival curve showing time to blood stage patency following intravenous injection of 1,000 salivary gland sporozoites, two biological replicates for clone 1, 1 replicate for wildtype and Clone 2, n = (wildtype = 5, Clone 1 = 10, Clone 2 = 5) I. qRT-PCR analysis of GAMA expression in *P. falciparum* NF54 infected mosquito midguts and salivary gland sporozoites, transcript abundance relative to 18s rRNA. RNA extracted from pools of >30 midguts or salivary glands for each timepoint.

**Figure 5:**
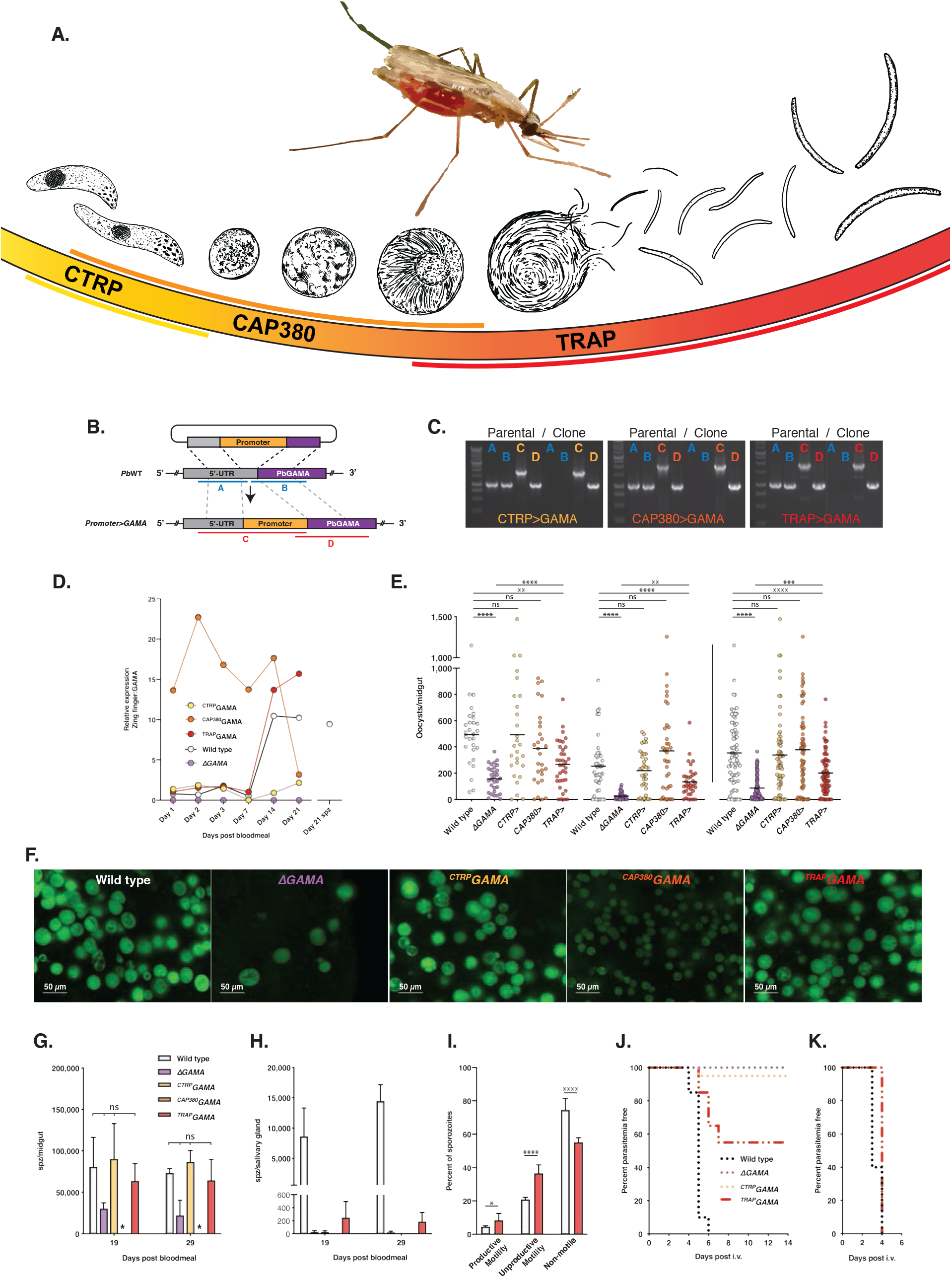
Parasites expressing GAMA under the control of stage specific promoters assist further characterization of protein function and suggest an essential role in hepatocyte invasion, but not merozoite egress from liver schizonts. A. Illustration of the expression profile desired from the promoter swap parasites. B. Schematic showing the method of genetic alteration used to generate parasites with GAMA expressed under different promoters, with approximate PCR amplicons indicated C. Electrophoresis gel of the genotyping PCR of mixed population of parental parasites following transfection (left of images) adjacent to a representative clone parasite for each of the promoter swaps (right of images), ^*CTRP*^*GAMA*, ^*CAP380*^*GAMA* and ^*TRAP*^*GAMA* (left to right). Approximate PCR amplicons annotated in (A). D. GAMA transcription levels based on qRT-PCR performed on cDNA prepared from pools of ∼50 midguts, per time point. Comparative C_T_ methods were used with primers specific for GAMA and PBANKA_0121700 (employed as a housekeeping gene). E. Day 14 oocyst counts with mean, two biological replicates, data to the right of the vertical line are pooled data, horizontal lines indicate mean (Mann-Whitney test, p > 0.05 = ns, p < 0.05 = *, p < 0.01 = **, p < 0.001 = ***, p < 0.0001 = ****). F. Live imagining of oocysts of each parasite line 21 days after infectious blood meal, wildtype, Δ*GAMA*, ^*CTRP*^*GAMA*, ^*CAP380*^*GAMA* and^*TRAP*^*GAMA* (left to right). G. Midgut sporozoites per mosquito at two timepoints, mean and SEM of two biological replicates. * ^*CAP380*^*GAMA* infected mosquitoes were dissected but no sporozoites detected. H. Salivary gland sporozoites per mosquito at two timepoints, mean and SEM of two biological replicates. I. Motility of wild type and ^*TRAP*^*GAMA* midgut sporozoites J. Kaplan Meier survival curve showing time to blood stage patency following intravenous injection of 200,000 midgut sporozoites per C57BL/6 mouse, four biological replicates combined, n = (wildtype = 20, Δ*GAMA* = 13, ^*CTRP*^*GAMA* = 20, ^*TRAP*^*GAMA* = 20). Twenty fields of a Giemsa-stained thin smear checked daily. K. Kaplan Meier survival curve showing time to blood stage patency following intravenous injection of 2,500 salivary gland sporozoites, n = 5 for wildtype and ^*TRAP*^*GAMA*.

Intensity of midgut infection was significantly lower for both *Pb*^*PfGAMA*^ clones across two of three biological replicates and remained significant when all data were combined (Fig 4C). However, the defect in midgut infectivity appeared less severe than for Δ*GAMA* parasites and when oocyst intensity as a percentage of wildtype control was calculated for Δ*GAMA* and *Pb*^*PfGAMA*^, there was a small but significant increase in the infection intensity for complemented parasites (unpaired t-test p = 0.041) (Fig. 4D). This partial restoration of ookinete midgut infectivity observed in *Pb*^*PfGAMA*^ parasites indicates a partial conservation of GAMA function between species.

Despite the greater number of oocysts in wildtype infected mosquitoes, there was no significant difference in the number of midgut sporozoites between wildtype and *Pb*^*PfGAMA*^ clones across multiple timepoints (Fig 4E). This could be due to the less pronounced reduction in oocyst numbers, but also the departure of large numbers of wildtype sporozoites from oocysts to the salivary glands that was not observed for *Pb*^*PfGAMA*^ parasites (Fig 4F) as well as greater nutritional stress within higher intensity wild type infections (39). Due to variation across biological replicates the difference in salivary gland numbers between wild type and *Pb*^*PfGAMA*^ parasites was not statistically significantly different, however the mean across all reps and timepoints were, 11,482 vs 188.1 and 85.2 spz for wild type, *Pb*^*PfGAMA*^C1/C2 respectively (if wild type sporozoite numbers are normalized to 100%, all p values are < 0.0001), and we therefore conclude that *P. falciparum* GAMA does not complement *P. berghei* GAMA function in this stage of the lifecycle. To investigate whether complementation had restored sporozoite egress from the midgut oocyst, the number of sporozoites in the salivary glands were converted to a percentage of the total collected (midgut and salivary glands combined), for wildtype, Δ*GAMA* and both complemented clones, in order to normalize for the slightly improved infections in *Pb*^*PfGAMA*^ infected mosquitoes. This revealed no improvement over Δ*GAMA* in the arrival of sporozoites to the salivary glands in complemented parasites (Supp. Fig 7A), and therefore complementation of *Pb*Δ*GAMA* parasites with *PfGAMA* does not overcome the sporozoite egress defect of Δ*GAMA*, indicating either a divergence of function at this stage or, an incompatibility of *Pb*^*PfGAMA*^ with interacting partners in *P. berghei* parasites.

To further investigate the role of *P. falciparum* GAMA in non-erythrocytic stages, 200,000 midgut sporozoites were injected intravenously into mice. Of 20 mice injected with *Pb*^*PfGAMA*^ sporozoites only 1 became patent (on day 6), in comparison with wild type, where all mice became patent by day 5 (Fig. 4G). *Pb*^*PfGAMA*^ was therefore unable to restore the infectivity of Δ*GAMA* midgut sporozoites to mice.

While the number of sporozoites detected in salivary gland dissections of *Pb*^*PfGAMA*^ was proportionally unchanged from Δ*GAMA* parasites (Supp Fig. 7), the increase in oocyst numbers meant that adequate salivary gland sporozoites were available to permit the assessment of their infectiousness to mice. All mice infected with 1,000 wild type salivary gland sporozoites became patent by day five, in contrast, only 3/15 mice infected with *Pb*^*PfGAMA*^ became patent on day 6 (Fig. 4H). Together these data indicate that the complementation of *PfGAMA* in Δ*GAMA P. berghei* parasites provide a partial rescue of the severe phenotype observed with Δ*GAMA*.

### A promoter swap strategy produces parasites with a range of GAMA expression profiles during mosquito development, permitting further investigation of protein function

A CRISPR/Cas9 methodology was utilized to insert ∼1 kb of the upstream promoter regions of stage specific genes into the *GAMA* locus in order to create parasites that express GAMA at specific timepoints during mosquito stage development. Selected promoters were CTRP, expressed only in ookinetes; CAP380, which is expressed throughout ookinete-oocyst development; and TRAP, which is expressed primarily in mature midgut sporozoites and salivary gland sporozoites (Fig. 5 A, Supp Fig. 8A) (40–44). To further confirm the temporal expression profile of these genes within infected mosquitoes in our insectary, midguts from wild type infected mosquitoes were collected at days 1-21 post infectious blood meal (d1 = ookinete, d2 = invaded ookinete transitioning to oocyst, d3 = very early oocyst, d7 = early-mid oocyst development, d14 = mid-late oocyst development and d21 = late-mature/egressed oocysts, salivary gland sporozoites were also collected at day 21). Quantitative RT-PCR was used to compare CT values (45) using primers targeting GAMA, the candidate promoter genes and PBANKA_0121700 as a housekeeping gene, revealing the expression profile of GAMA and also the candidate promoter genes. These qPCR data demonstrated an accurate match with previously published data (40)(our selection of PBANKA_0121700 over more commonly used housekeeping genes is detailed in Supp Fig. 8A-I) and was consistent with our previous observations of the expression profile of GAMA in our GAMA-mCherry tagged parasites (Supp. Fig. 5). *CTRP, CAP380* and *TRAP* revealed the anticipated expression profiles (Supp Fig. 8I). The parasites ^*CTRP*^*GAMA*, ^*CAP380*^*GAMA* and ^*TRAP*^*GAMA* were consequently generated and used for further analysis (Fig. 5B & C).

In order to confirm that the promoter swaps had produced the desired expression profiles, midguts infected with each parasite line were again collected at days 1, 2, 3, 7, 14 and 21 post infectious blood meal and GAMA transcript abundance determined by qRT-PCR. The expression profiles of GAMA under these chosen promoters correlated well with anticipated expression, albeit at lower levels than for their own respective transcripts. This was however fortuitous, as this reduction brought expression of GAMA under the promoter swaps to similar intensities as the endogenous expression of GAMA, which is itself relatively low (Fig. 5D). We therefore used these parasites to further investigate the role of GAMA in mosquito stages.

### Promoter swap parasites confirm essential functions of GAMA across parasite mosquito stages

#### ^CTRP^GAMA

Ookinetes of these parasites were fully infectious to mosquito midguts and oocyst counts were no different to wild type, further confirming the role of GAMA in midgut invasion (Fig 5E). This functional maintenance was predicted from the ookinete stage expression profile of CTRP (Fig. 5A). Oocysts appeared morphologically indistinguishable from wildtype (Fig. 5F) and generated similar numbers of sporozoites (Fig. 5G). Expression data revealed low level GAMA expression late in oocyst development in ^*CTRP*^*GAMA* parasites, a phenotype previously documented in a study which used the *P. berghei CTRP* promoter region for GFP expression (46); however, sporozoites were not found in significant numbers in the salivary glands (Fig. 5H), indicating inadequate expression/translation of GAMA to rescue oocyst egress. To investigate infectivity, 200,000 midgut sporozoites were injected intravenously into mice, where all but one remained uninfected (Fig. 5J), likely due to the very low-level GAMA. This further showed that GAMA has an important role in ookinete infection of the mosquito vector, egress of sporozoites from the oocyst and infection of the mammalian host.

#### ^CAP380^GAMA

^*CAP380*^*GAMA* parasites were as infectious to mosquitoes as wild type, in keeping with the GAMA transcript abundance found in ^*CAP380*^*GAMA* ookinetes (Fig. 5D & E). ^*CAP380*^*GAMA* oocysts however exhibited a striking developmental defect, whereby oocysts exhibited lower level GFP expression that wild type and did not grow beyond ∼day-five size (Fig 5F), indicating that high GAMA expression during oocyst development, which is turned off in wild type parasites at this stage, has a detrimental impact on oocyst development. In keeping with the minimal oocyst growth, no sporozoites were formed in these oocysts (Fig. 5G & H) and therefore investigating the invasion phenotype of GAMA negative salivary gland sporozoites, the intended objective for this parasite, was not possible.

#### ^TRAP^GAMA

^*TRAP*^*GAMA* parasites displayed an intermediate midgut infectivity phenotype, being both significantly less infectious than wildtype (mean oocyst/midgut of two biological replicates pooled: wild type = 354, ^*TRAP*^*GAMA* = 201, p = 0.0002), and significantly more infectious than Δ*GAMA* (Δ*GAMA =* 87 oocysts/midgut, p = <0.0001). TRAP is expressed at very low level in ookinete stages (Supp Fig. 8I), and in the ^*TRAP*^*GAMA* parasites, low-level GAMA transcription was also observed (Fig 5D), thereby potentially explaining the intermediate midgut infectivity phenotype (Fig. 5E).

Oocysts were morphologically indistinguishable from wildtype and developed normal numbers of sporozoites (Fig. 5G). However, contrary to our expectations of functional oocyst egress based on equivalent GAMA transcript abundance (Fig. 5D), these sporozoites were unable to exit oocysts and were only recovered in small numbers from salivary gland dissections (Fig. 5H). Given the similarity in expression level of GAMA between wild type and ^*TRAP*^*GAMA* parasites an oocyst egress defect was unexpected. Sporozoite motility has been shown to occur prior to oocyst egress (28) and with the knowledge that Δ*GAMA* oocysts showed a motility defect (Fig. 1I), the motility of ^*TRAP*^*GAMA* sporozoites was investigated. Surprisingly, motility data for ^*TRAP*^*GAMA* midgut sporozoites revealed these parasites to be more motile in both categories of motility than wildtype (productive motility p = 0.0136, unproductive p < 0.0001) (Fig. 5I), perhaps due to a build-up of mature, motility-competent-yet-egress-defective sporozoites when compared to egress-competent wild type. The unusual split in the phenotype of ^*TRAP*^*GAMA* parasites, *i*.*e*., egress-deficient but motility-competent, suggests a dual functionality of GAMA, both in motility and an unknown, motility-independent aspect of oocyst egress. When mice were injected with 200,000 ^*TRAP*^*GAMA* midgut sporozoites, 9/20 mice became patent whereas all mice infected with wild type became patent by day 6 (Fig 5J), indicating that GAMA plays a role in mammalian infection independent of motility. To explore the phenotype further, a limited number of sporozoites were obtained from salivary gland dissections to investigate their infectivity to mice. When injected with 2,500 wildtype or ^*TRAP*^*GAMA* salivary gland sporozoites, no difference was observed in the delay to blood stage patency, revealing that ^*TRAP*^*GAMA* salivary gland sporozoites were as infectious to mice as wildtype (Fig. 5K). A recently published *P. berghei* merosome proteome failed to detect TRAP (47). Therefore, the ^*TRAP*^*GAMA* parasite is unlikely to express GAMA during liver stage development. These data, combined with the well-defined sporozoite-stage function of TRAP (37,44,48,49), indicate there would be an absence of GAMA in the merosomes of ^*TRAP*^*GAMA* parasites and therefore further support a role for GAMA during the initial infection of the vertebrate host, not during the egress of merozoites from hepatocytes.

## DISCUSSION

Micronemes, rhoptries and dense granules (listed in order of their release) are the invasive organelles which store and discharge effector molecules that mediate critical aspects of parasite/vector and parasite/vertebrate host interactions. Of these, only micronemes are present in all zoite forms. The effector molecules released by micronemes can have broad roles in parasite life cycle progression or can play rather narrow roles in a particular zoite activity. In this current study our results demonstrate that GAMA exerts critical functions at multiple stages of the *Plasmodium* life cycle, including the productive invasion of the mosquito midgut by ookinetes, the egress of sporozoites from the mature oocyst, the gliding motility of sporozoites and their infection of the vertebrate liver. The breadth of involvement of GAMA in the microneme associated functions of egress, motility and invasion, across multiple life cycle stages point to an auxiliary role in the basic functioning of micronemes.

Ookinetes must traverse through the injested blood meal, peritrophic matrix and epithelial cell layer of the mosquito midgut to access their destination niche at the basal lamina of the mosquito midgut. For these functions, ookinetes contain many micronemes, laden with proteins required for this journey, many of which with direct roles in either motility (eg. CTRP (41)), or enzymatic functions (eg. chitinase (50)), to clear a path through this environment. Parasites with GAMA expression under the CTRP and CAP380 promoters, both active in ookinetes were equally as infectious as wild type parasites, further confirming a role for GAMA in the midgut invasion process.

RT-qPCR data revealed GAMA transcripts elevated towards the end of oocysts development. mCherry tagged parasites further confirmed the tight temporal expression of GAMA only within oocysts containing well segregated sporozoites, in keeping with observations during the blood stage of *P. falciparum* where GAMA is only expressed late in schizogony (13). Micronemes contain proteins which possess aggressive qualities, such as digesting extracellular matrices (Chitinase (52)) and for punching holes in membranes (CelTOS (51)), to access the basal lamina of the midgut and it therefore makes sense that micronemal proteins would only be expressed at a time when their functions are essential for life cycle progression. The ^CAP380^GAMA parasites which continually expressed GAMA from ookinete stages onwards and throughout oocyst development, exhibited developmental dysregulation, confirming the importance of the timing and perhaps the correct packaging of GAMA within micronemes, which after oocyst development is initiated, is then only expressed late in oocyst development (8) to allow for sporozoite egress from the mature oocyst.

Haemolymph collection confirmed an essential role of GAMA in sporozoite egress from oocysts. Motility prior to egress has been reported for oocyst sporozoites (28) and therefore the egress defect observed in Δ*GAMA* parasites could be attributed to the motility defect we observed in midgut sporozoites. However, the unusual specificity of defects in the ^*TRAP*^*GAMA* parasite excludes motility as the likely cause of the egress defect, as the oocyst sporozoites of this parasite line exhibited *increased* motility compared to wild type controls but were still unable to egress. In wild type parasites, the release of TRAP from micronemes and consequent initiation of motility is likely accompanied by the release of other classes of micronemes with egress functional cargo. Electron microscopy has previously revealed uneven distribution of proteins between micronemes (53–56), and in *T. gondii* an investigation into the trafficking of micronemal proteins revealed distinct subsets of micronemes and egress and invasion mutants exhibited atypical trafficking of specific micronemal proteins (57). Therefore, individual micronemal populations likely perform specific and exclusive roles (58,59). While GAMA transcripts produced under the *TRAP* promoter were found at a similar abundance to those of wildtype parasites, the *TRAP* promoter may have resulted in GAMA localization specifically within subsets of motility specific micronemes normally occupied by TRAP and not in all microneme populations which would otherwise contain GAMA. It is possible that the localisation of GAMA under the TRAP promoter was atypical, resulting in GAMA localization within only TRAP containing micronemes, and not those responsible for egress functionality.

While productive motility was absent from Δ*GAMA* sporozoites, they were capable of patch gliding, phenocopying parasites with mutations or truncations of the cytoplasmic tail of TRAP (32,60) or with defects in their ability to cleave TRAP from their exterior surface (37) and indeed Δ*TRAP* parasites (29). The similarity in motility phenotype between Δ*TRAP* and Δ*GAMA* parasites suggests that TRAP function is somehow reliant on the presence of GAMA. As has been previously suggested (28), it is likely that GAMA is required for the correct functioning of micronemes and positioning of proteins on the parasite surface where effectors of motility (29,44,61,62), traversal and invasion (51,53,63–67), perform their functions.

GAMA was found to be shed in the trails of gliding salivary gland sporozoites and joins three other parasite proteins previously detected in trails; TRAP, CSP, and CelTOS (30,37,54). mCherry was not readily detected on the surface of *GAMA-mCherry* sporozoites by IFA, perhaps due to the position of the mCherry tag close to the GPI attachment which may shield it from antibody access as has been shown for other sporozoite surface proteins, such as SSP3 (38). Alternatively, it may only be present in small amounts. Indeed, TRAP, which is expressed at higher levels than GAMA (Supp. Fig. 8I) (40,68), is also not easily detected on the surface of sporozoites by IFA (53). In addition to the internal micronemal localization, endogenous mCherry in live, motile *GAMA-mCherry* sporozoites revealed, prior to photobleaching, an increase in mCherry at the sporozoite periphery, again consistent with the observed deposition in trails.

GAMA shedding during sporozoite motility is in line with previous observations, where GAMA was found to be shed from the surface of *P. falciparum* merozoites during red blood cell invasion by a protease other than SUB2 (13). The shedding of GAMA in this instance was attributed to invasion-associated shedding, akin to that observed with MSP1 and AMA1 (69–71). Merozoite motility was recently characterised for the first time in both *P. falciparum* and *P. knowlesi* (72) and a conserved role of GAMA across zoite stages would implicate the involvement of GAMA in both motility and invasion of merozoites.

Despite expression across all invasive life cycle stages, GAMA has remained understudied and has not been widely recognized in literature concerning proteins essential for merozoite, ookinete and sporozoite function. Considering the broad involvement across all processes associated with micronemes, we propose a model whereby GAMA is required for microneme function and by proxy the phenotypes associated with micronemal proteins at large. The signaling pathways that lead to micronemal release and the associated parasite activity, while complex, are being unraveled. The essential nature of all the processes in which GAMA is involved warrants further investigation of this key parasite protein.

## METHODS AND MATERIALS

### Bioinformatics analysis

DNA and protein sequences were retrieved from plasmodb (https://plasmodb.org/plasmo/) with sequence alignments performed in Geneious version 10.2.3. Protein features were predicted with the following online tools: Signal peptides with SignalP-5.0 (http://www.cbs.dtu.dk/services/SignalP/), transmembrane domains with TMHMM Server v. 2.0 (http://www.cbs.dtu.dk/services/TMHMM/), GPI anchors with (http://gpcr.biocomp.unibo.it/predgpi/pred.htm), molecular weight (https://www.bioinformatics.org/sms/prot_mw.html) and the presence of other protein domains were predicted with InterPro (https://www.ebi.ac.uk/interpro/).

### Global diversity of *gama*

The MalariaGEN Pf3K project release 5.1 data (73) was used to estimate global diversity of *gama*. The publicly available dataset includes whole genome sequencing data from 2,512 samples collected in multiple locations in Asia and Africa in addition to 156 additional isolates from Papua New Guinea (74). Briefly, GATKv4.0 was used for variant calling and samples containing low quality bases were removed. Detailed bioinformatics processes can be found at https://github.com/myonaung/Pf3K_Scripts. Singleton SNPs were converted back to reference to prevent false positive variants, and only major allele was included in the analysis for mixed infection samples(75). R software, VaxPack (https://github.com/BarryLab01/vaxpack) was used for global population genetic analysis.

### Genetic modification of *P. berghei*

To generate the knock-out and *P. falciparum GAMA* knock-in parasite lines the Gene In Marker Out (GIMO) methodology was employed, as previously described (25). In brief, the GIMO plasmid was modified with the addition of 5’ and 3’ homology arms corresponding with the 5’ and 3’ UTR of GAMA, flush to the ORF of GAMA, so that a KO parasite would be generated expressing the hdhfr/yfcu cassette in place of the GAMA gene. Transfections were performed as previously described (76), on the marker free, genome integrated GFPLuc expressing *P. berghei* parasite line Pb676, which was used as wild type throughout this study {Janse:2006du}. Transfectant parasites obtained after negative selection with pyrimethamine were cloned by limiting dilution into naïve recipient mice to obtain clonal parasites (genotyping primer sequences in Supplementary).

To generate the *Pb*^*PfGAMA*^ parasite line, the full-length *P. falciparum* GAMA gene was amplified from *P. falciparum* genomic DNA and inserted via the KpnI and SacII restriction sites into the *PbGAMA* knockout GIMO plasmid (Supplementary primer table). Transfections were performed on Δ*GAMA* parasites and integration of *PfGAMA* was achieved by negative selection with 5-fluorocytosine, administered *ad libitum* in the drinking water of the mice at a concentration of 1 mg/ml. Clonal parasites were obtained as above.

### Generation of promoter switch and Cherry tag parasite lines

A CRISPR/Cas9 methodology was used to generate tagged and promoter switch parasites. To generate promoter switch parasites, guides were identified in the 188 bp region just upstream of the GAMA ORF. Homology arms corresponding to the 758 bp upstream of this region (5’ flank), or the first 768 bp of the GAMA gene (3’ flank) were amplified from genomic DNA and ligated into the pYC-L2 plasmid (PMID: 24987097) via restriction cloning (primers used for all plasmid construction and genotyping are listed in Supplementary Table 1). Stage specific promoters were amplified from genomic DNA and ligated between the homology arms; CTRP = 781bp, CAP380 = 1065, TRAP = 946.

For mCherry tagging, homology arms for the tagging construct were designed so that the tag would be inserted between bases 1818-1819 of the ORF, corresponding to an insertion 19 aa from the end of the GAMA protein, 5 residues downstream of the predicted GPI anchoring site. Four guides were designed close to the intended insertion site of the tag with multiple shield mutations built into the repair construct (primer table). Of these guides, guides 9 and 42 were used in combination. Repair constructs were synthesized by Genscript and ligated into the pYC plasmid by restriction cloning with KpnI and NotI.

Single stranded guide primers were annealed in a 50 µL volume (10 µL of each 100 µM primer with 25 µL water and 5 µL cutsmart buffer), by heating to 95 ºC for 5 minutes and allowing to cool to room temperature. Annealed guides were inserted into the pYC-L2 plasmid, via restriction cloning with the enzyme Esp3I. Guide integration was confirmed via sanger sequencing with the primer CJ059.

Transfection and downstream cloning of transgenic parasites was performed as detailed above. The GAMA ORF was Sanger sequenced to ensure fidelity.

### Mosquito infections

*Anopheles stephensi* were fed on *P. berghei* infected Swiss Webster mice (Envigo) following standard methods described in (77). Mosquitoes were maintained at 19 ºC with a 12/12 day/night lighting pattern.

### Ookinete preparation and counting

22 hours after infectious blood meal the midguts of ∼20 mosquitoes were isolated into one well of a staining dish containing 200 µL PBS. Engorged midguts were pipetted up and down to homogenise contents until the suspension was smooth, then transferred to a 1.5 mL Eppendorf tube and centrifuged for three minutes at 800g. The supernatant was removed, and the pellet resuspended in 1 mL of 0.15% saponin in PBS. This was incubated for 10 minutes on ice, agitating frequently by gentle vortexing. After saponin lysis the tube was centrifuged for 3 minutes at 800g and the supernatant removed and the pellet washed with 1 mL of PBS, three times. The ookinete preparation was resuspended in a known volume of PBS and spotted onto slides for Giemsa staining and counting.

### Oocyst counts

All parasites used in this study were derived from Pb676, a marker free GFP reporter line and therefore oocyst numbers could accurately be determined by fluorescence microscopy. Midguts were removed from mosquitoes 14 days post infectious blood meal and the green channel imaged on a Keyence microscope under 100 or 40x magnification. Oocyst numbers were counted manually in FIJI using the cell counter plugin. To compare oocyst numbers between parasite lines, nonparametric statistical analyses were performed to calculate differences between median oocyst intensities (Mann Whitney test [two tailed with additional Mann-Whitney CI]).

### Sporozoite counts

Mosquitoes were aspirated into 70% ethanol and subsequently rinsed in PBS prior to dissection. Where midgut and salivary gland sporozoites were collected from the same mosquitoes, salivary glands were collected first, then midguts. Salivary glands were isolated using the “dirty” method of dissection, whereby mosquito heads were cut off with a sharp 23-gauge syringe needle and the thorax depressed to release salivary glands from the thorax which were consequently collected in Schneider’s medium with a fine glass Pasteur pipette. Midguts were extracted with fine forceps and collected into a 1.7 mL Eppendorf tube containing Schneider’s medium, with care taken to avoid damaging the midguts during dissection. Salivary glands or midguts were centrifuged at 8000 g for 1 minute and then crushed with a plastic pestle, thrice, prior to dilution and quantification on a haemocytometer. In all instances >25 mosquitoes were dissected and pooled together for sporozoite quantification.

To collect haemolymph the last two segments of the abdomen were removed with a 23-gauge needle, a 31-gauge needle containing Schneider’s medium was then inserted into the top of the thorax and 15 drops of Schneider’s medium passed through each mosquito and collected. This was concentrated and sporozoite numbers enumerated with a haemocytometer. The exsanguinated mosquitoes were then dissected for their midguts which were processed for sporozoite counting as detailed above.

### Imaging

Schizonts, ookinetes and sporozoites, having been fixed in PFA were permeabilised in 0.5% TRITON X100 for 15 minutes at RT. Blocked for 30 minutes at 37 °C in 3% BSA in PBS. Incubated with primary antibodies; rat anti mCherry monoclonal 16D7 (Thermo Fisher) 1:500, Rabbit anti Py-MTIP 1:1000 (PMID: 12456714) at 37°C for 45 minutes. After 3 washes in PBS, samples were incubated with appropriate secondary antibodies at 1:1000 and DAPI 1µm/mL in 3% BSA for 30 minutes at 37 °C. Samples were again washed and mounted in Fluoromount G, stored at RT overnight and imaged the following day on a DeltaVision microscope.

Oocysts of various maturity were imaged on a Keyence or DeltaVision microscope using the endogenous GFP and mCherry fluorescence with DAPI staining. The same method was used for imaging endogenous GFP and mCherry fluorescence in sporozoites. Line plot pixel intensity analysis was performed in FIJI.

### Sporozoite infections of mice

7-10 week-old C57B/6 mice were anaesthetized in an isofluorane chamber and injected retro-orbitally with the required number of sporozoites in a 200 µL volume, for accuracy. Thin smears were prepared daily from day two onward and 20 fields per slide examined for the presence of blood stage parasites. All sporozoite derived blood stage infections from non-wild type parasites used in this project were genotyped to confirm the transgenic nature of the blood stage parasites.

### Motility assays

Salivary gland sporozoites were allowed to glide on cover slips in Schneider’s medium containing 1% BSA and imaged at 1000x on a DeltaVision microscope recording both Brightfield and TRITC channel images every 2 seconds.

Midgut sporozoites were purified from midguts on an Accudenz gradient as previously described (78). The interface was recovered with a glass Pasteur pipette and made up to 2 mL with Schneider’s media and spun at 8000g for 3 minutes to pellet sporozoites. Sporozoites were washed once with Schneider’s medium and the supernatant aspirated leaving 20-30µL volume. 2.5 µL of sporozoite suspension was mixed with 0.5 µL of 9% BSA and 3 µL of Matrigel. The mixture was thoroughly pipetted up and down and spotted onto a slide. An 18 mm cover slip with Vaseline buffer spots, to prevent total flattening of the preparation was carefully placed over the mixture, sealed with nail varnish and left to set at 37 c for 7 minutes, upside down. Motility time-lapse movies were recorded on a Keyence microscope at 200x magnification, 1 fps for 7 minutes.

Motility trails. 8 chamber slides (LabTek) were coated with FBS for 1 hour at 37 c, FBS removed and washed once with PBS. 10,000 wild type or GAMA-mCherry salivary gland, Accudenz purified, sporozoites in RPMI with 1% BSA were added per well, spun down at 50 g for 3 minutes and kept at 37 c. After 45 minutes, media was removed, and wells fixed with 4% PFA. To avoid cross reactivity between closed related antibody species, rat and mouse, trails were stained first with the aforementioned anti mCherry antibody, followed by appropriate 594 secondary. The slide was again fixed with 4% PFA and then stained with a 647 conjugated anti CSP antibody 3D11. After washes slides were kept in PBS and imaged under 1000x magnification on a Keyence microscope. Imaging settings were maintained consistent between mCherry and wild type sporozoite wells.

### Quantitative PCR

Midguts were collected from mosquitoes on days 1, 2, 3, 7, 14 and 21 post *P. berghei* blood meal into PBS on ice. For wild type infected mosquitoes, salivary gland sporozoites were also collected on day 21. Midguts of NF54 infected mosquitoes were collected on days 2, 3, 6, 10 and 14 post bloodmeal and salivary gland sporozoites also collected on day 14. Supernatant was removed from midguts and sporozoites after spinning at 8000g and 16,000g respectively and they were snap frozen and stored on dry ice. RNA was extracted with a Qiagen RNeasy kit. A DNase digestion step was performed on the column to remove any contaminant gDNA with the Qiagen DNase kit. RNA was confirmed negative for gDNA by PCR prior to preparation of cDNA with the SuperScript VILO cDNA synthesis kit using 2 µg of RNA per reaction. cDNA was diluted 1:50 in water prior to qPCR with SYBRgreen (biomake) on the QuantStudio3 platform, using the comparative *C*_T_ method for data analysis (45).

## ETHICS STATEMENT

The study was performed in strict accordance with the recommendations in the Guide for the Care and Use of Laboratory Animals of the National Institutes of Health (NIH), USA. To this end, the Seattle Children’s Research Institute (SCRI) has an Assurance from the Public Health Service (PHS) through the Office of Laboratory Animal Welfare (OLAW) for work approved by its Institutional Animal Care and Use Committee (IACUC). All of the work carried out in this study was specifically reviewed and approved by the SCRI IACUC.

## ACKNOWLEDGEMENTS

We would like to thank the dedication of the Research Insectary for mosquito production. Research reported in this publication was supported by the National Institutes of Health under awards R21AI146391 (AMV), K25AI119229 (KES), R01AI148489 (KES) and R01GM087221 (RLM), the PATH Malaria Vaccine Initiative, and by internal funds from the Center for Global Infectious Disease, Seattle Children’s Research Institute (AMV). The content is solely the responsibility of the authors.

